# Presence of *Triatoma breyeri* (Reduviidae, Triatominae) in Bolivia

**DOI:** 10.1101/2024.03.18.585491

**Authors:** Frédéric Lardeux, Alberto Llanos, Roberto Rodriguez, Luc Abate, Philippe Boussès, Rosenka Tejerina Lardeux, Christian Barnabé, Lineth Garcia

## Abstract

The study focuses on identifying and understanding the ecological dynamics of *Triatoma breyeri* in Bolivia. Morphological identification and molecular analysis using gene fragments (*COI, CytB* and *16S*) confirms *T. breyeri’s* presence and its relation to other species. The species has been consistently found in the Estancia-Mataral – La Palma region since 2010 but has not spread to other regions in Bolivia. The region of occurrence is a small characteristic dry inter-Andean valley. A MaxEnt model suggests part of the Bolivian Montane Dry Forest ecoregion serves as a unique habitat within its range. The infrequent presence in Bolivia and the distance from its main range in Argentina suggest recent accidental introduction, possibly through human transport. Further research is needed to comprehend its persistence in this small area of Bolivia.

## Introduction

In the realm of entomology, it is not uncommon to encounter instances where a particular insect species manifests its presence far beyond the confines of its established geographic distribution range. This phenomenon can be driven by a multitude of interconnected factors, including climate change, habitat alterations, human-mediated dispersal, and natural range shifts. The typical range of a species often emerges as a product of extensive evolutionary processes and adaptations finely tuned to the specific environmental conditions prevailing within it. However, there are occasions when species are discovered in locations far removed from their conventional geographic boundaries. One intriguing facet of these expansions involves the transportation of species to entirely new areas, where they not only survive but also establish self-sustaining populations. This phenomenon underscores the remarkable adaptability and resilience of insect species in the face of ongoing environmental changes.

In this context, our study explores a recent discovery: the presence of the triatomine bug *Triatoma breyeri*, Del Ponte 1929, well beyond its customary geographic range in Argentina.

The discovery of *T. breyeri* in a small valley of southern Bolivia occurred serendipitously in 2010, during samplings conducted within the scope of an investigation into insecticide resistance in another triatomine bug, *Triatoma infestans* (Klug 1834), the main vector of Chagas disease in this country. Remarkably, years later, local residents also brought spontaneously additional specimens of *T. breyeri* to local health centers, and authors collected few specimens, further confirming its presence in the area.

In this study, a brief overview of *T. breyeri* is provided, including its characteristics and its usual distribution range, before delving into the circumstances surrounding its unforeseen discovery in Bolivia, elucidating the mechanisms that may have underpinned its geographical expansion. A Maxent model of habitat suitability is developed to gain deeper insights into its dispersal dynamics in Argentina and Bolivia.

## Results

### Concise overview of knowledge on *T. breyeri*

The initial specimens of *T. breyeri* were collected in 1929 within the province of La Rioja, Argentina, which subsequently served as the foundation for the species’ formal description [1]. During the 1950s, the species remained relatively understudied, despite the collection of numerous specimens in that region. Indeed, at that time, its distribution was primarily constrained to the provinces of Catamarca and La Rioja in Argentina [2]. Subsequently, the species was also documented in the province of Córdoba [3] and within the province of Santiago del Estero, near its borders with Tucumán and Catamarca, as supported by a collection carried out in 1977 (GBIF data).

*T. breyeri* can be readily identified by examining the morphological characteristics of both adults and larval stages [2–4] and even its egg has been described [5].

The taxonomic classification of *T. breyeri* remains a subject of ongoing debate within the scientific community. Initially, it was assigned to the “*spinolai*” complex, a group introduced in 1979 by Lent and Wygodzinsky to describe “*a small, taxonomically isolated, and geographically limited group inhabiting semiarid regions of southern South America. This complex encompassed three species: T. spinolai (found in central and northern Chile), as well as the central and western Argentinian species T. eratyrusiformis and T. breyeri*” [3].

Subsequently, with the establishment in 1940 of the *Mepraia* genus, *T. spinolai* Porter 1934 was reclassified as *M. spinolai*. Moreover, further complicating matters, *T. breyeri* was eventually placed within the *Mepraia* genus in 2002 as *M. breyeri* [6]. The *spinolai* complex became the *Mepraia* complex where all the species (the 3 above mentioned species plus *M. gajardoi* Frias, Henry & Gonzalez, 1998 and *M. parapatrica* Frias, 2010) constitute a monophyletic group [6–8]. However, some authors maintain a “*breyeri*” complex, which comprises of the only two species from the eastern sides of the Andes: *T. breyeri* and *T. eratyrusiformis* [9, 10] while the *Mepraia s.s*. species from the western side of the Andes, are maintained in the *Mepraia* complex. It is important to note that these taxonomic rearrangements remain contentious and continue to be a subject of dispute in the scientific community [11, 12]. In La Rioja Province, a subspecies known as *T. breyeri dallasi* was described [13]. Although it has not undergone formal synonymization [3], some authors have proposed such synonymy [2, 14, 15]. Nevertheless, the validation of this subspecies continues to be a subject of debate [16].

The ecosystem where *T. breyeri* grows has been described as a salty semidesert ecosystem around Salinas Grandes in central Argentina, ranging from 200 to 700 meters above sea level [9]. As of the present day, as revealed by the 12 distinct collecting points from the GBIF database (https://www.gbif.org/), the species’ geographic range is acknowledged to be restricted to the Andean pre-cordillera within these Argentinian provinces. These occurrences fall inside the two ecoregions Dry Chaco and High Monte as defined by the WWF (World Wide Fund for Nature) [17] (Fig. 1). The Dry Chaco ecoregion is primarily situated in the northwestern two-thirds of western Paraguay, extending eastward across the Andes into southeastern Bolivia and northwestern Argentina. The climate is dry, with an annual rainfall of 650-350 mm, and an average temperature of 28°C-12°C. The Chaco region comprises various habitats, including savannas, thorn forests, and transitional areas between these two dominant types. The High Monte ecoregion is situated along the eastern slopes of the Andes of Argentina, stretching from the province of Salta in the north to the province of Mendoza in the south. This landscape is characterized by rugged mountains and enclosed basins nestled between the Sierras Pampeanas to the east and the primary Andean mountain range to the west. The climate in this region is characterized by temperate, arid, and semi-arid conditions. Annual precipitation levels in the High Monte ecoregion vary depending on geographic location and elevation, ranging from under 100 mm to approximately 450 mm [18]. More precisely, the *T. breyeri* occurrences from the GBIF data base lie within several Köppen-Geiger climate classification [19]: BWh (arid, desert, hot) [2 collecting points], BWk (arid, desert, cold) [2 collecting points], BSh (arid, steppe, hot) [5 collecting points], Cwa (temperate, dry winter, hot summer) [2 collecting points], and Cwb (temperate, dry winter, warm summer) [1 collecting point] (Fig. 2).

**Fig. 1.**
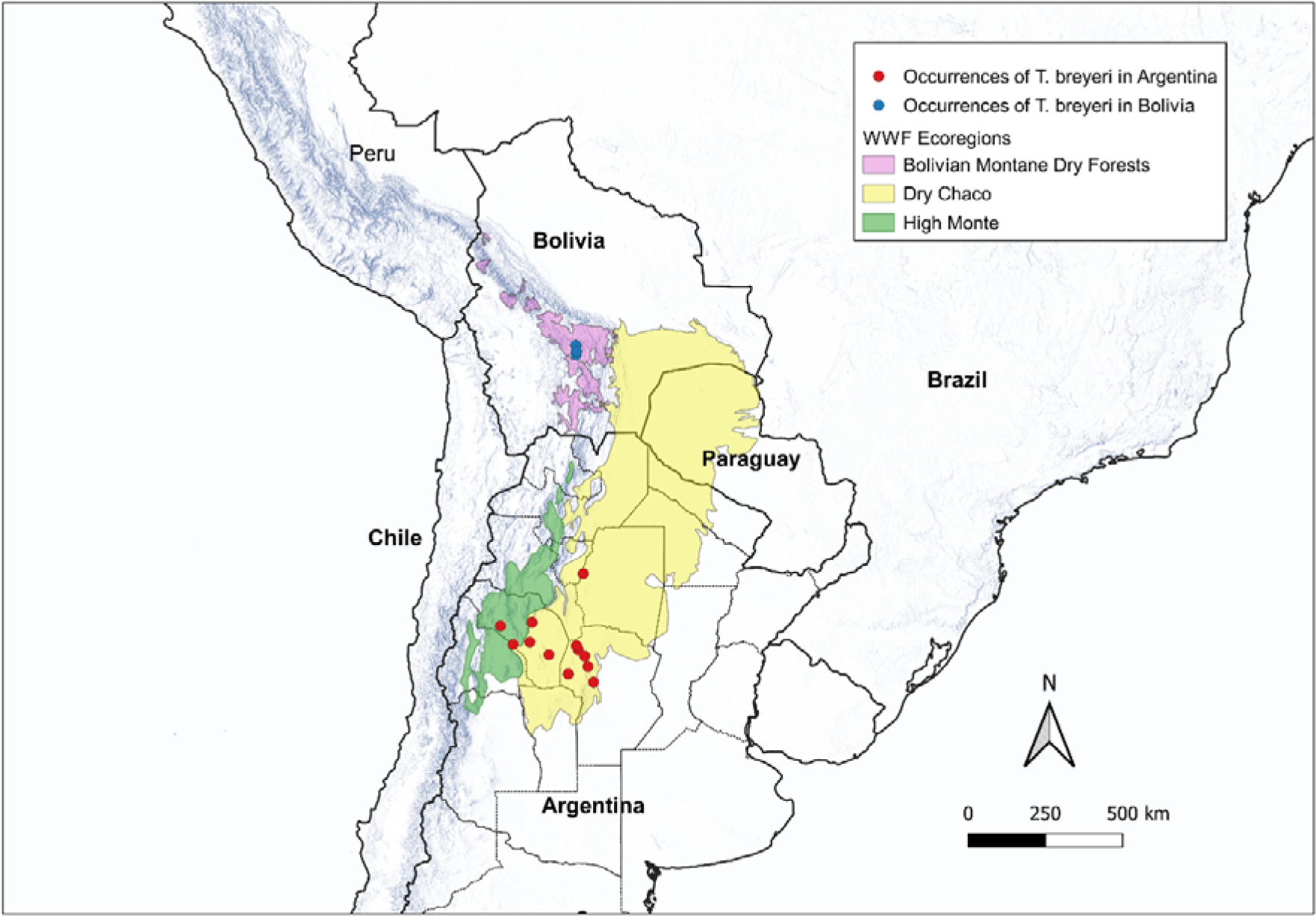
Collecting points of *T. breyeri* in Argentina and Bolivia and their related WWF ecoregions.

**Fig. 2.**
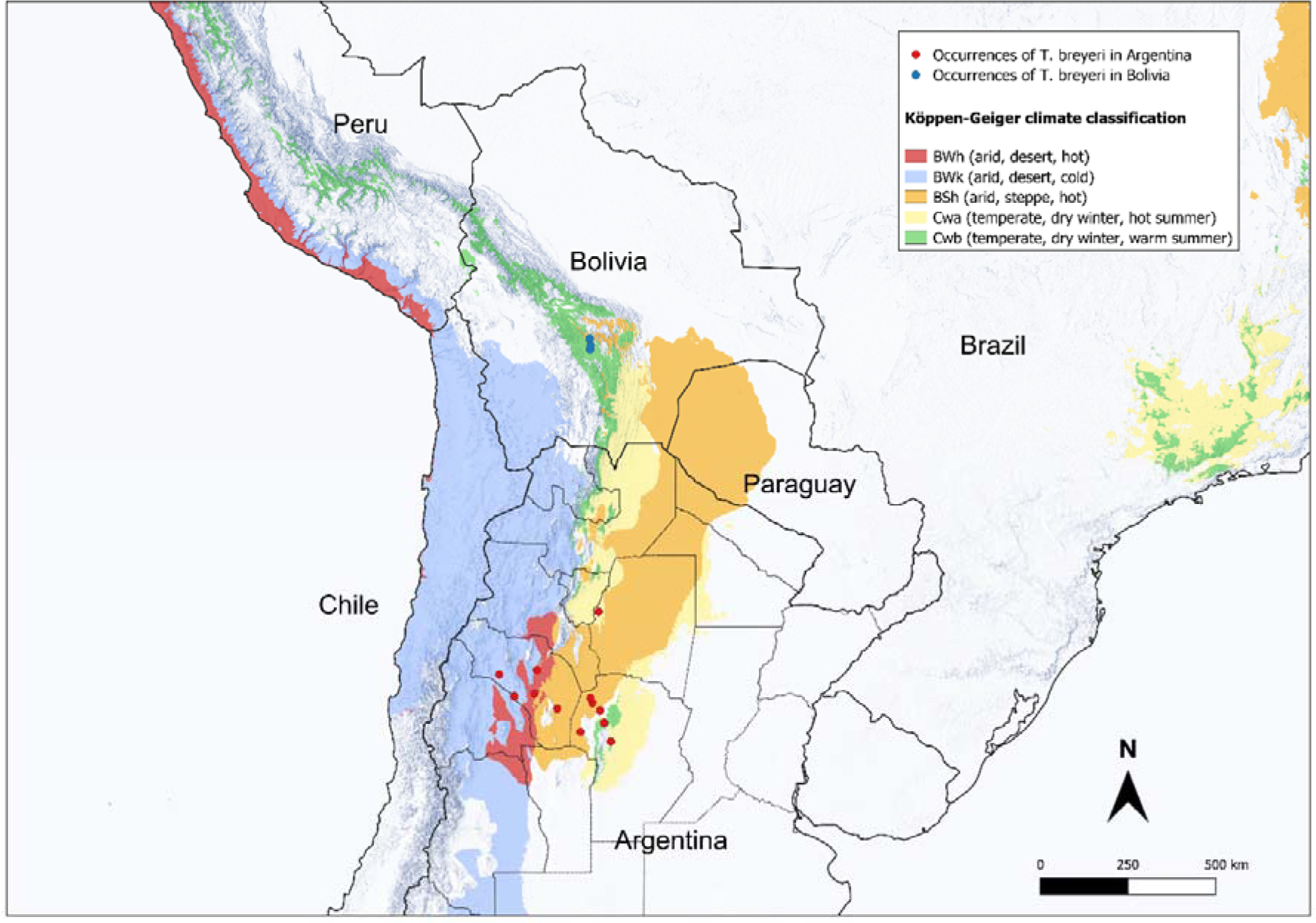
Collecting points of *T. breyeri* in Argentina and Bolivia and their related Köppen-Geiger climate classes.

The ecological characteristics of *T. breyeri* are only partially understood. It predominantly inhabits the lower sections of branch fences, typically in proximity to the burrows of small rodents of the Caviidae and Cricetidae family [3]. It is reported that during the summer, adults can be encountered in the vicinity of houses and, occasionally, inside rooms [20–22]. However, it is important to note that they do not establish colonies within domestic environments [21, 23, 24]. Laboratory experiments conducted at a controlled temperature of 25°C (±4°C) and relative humidity of 60% (±20%) have revealed that *T. breyeri* exhibits some resistance to fasting and demonstrates a low population growth rate (*r* = 0.017) with limited dispersal capacity. The nymphal stage spans approximately 318 days (comprising 37 days for N1, 35 days for N2, 56 days for N3, 87 days for N4, and 103 days for N5). Adult individuals have an average lifespan of ≈20 days, and the incubation period for eggs lasts ≈ 32 days. The generation time is estimated to be approximately 58 days (*s.d*. ≈ 4), while the *R_0_* value is calculated to be 2.68 (*s.d.* ≈0.3) [25]. The species is not known as a vector of *Trypanososma cruzi*, the causative agent of Chagas disease [3], but may carry the entomopathogenous virus *Triatoma virus* [26].

### Discovery in Bolivia

In Bolivia, specimens of *T. breyeri* were first collected in 2010, in the small locality of Estancia-Mataral (lat. −18.5974, lon. −65.1463, alt. 1520 m), within a peri-domestic environment. Its discovery occurred during entomological surveys (*i.e.*, active search in households by specialized technicians) conducted by the National Chagas Program of Bolivia as part of the efforts to control *T. infestans* the main vector of Chagas disease in Bolivia. Estancia-Mataral has no more than 60 households and lies where the small river Rio Novillero joins the Rio Grande, approximately 90 km by road, north of Sucre, the country’s administrative capital. Live specimens were transferred to the entomology laboratory at the Escuela Técnica de Salud Boliviano Japonesa de Cooperación Andina in Cochabamba, where they were reared under controlled conditions (28°C, 70% relative humidity), with monthly feeding on hens, for several years by one of us (RR). More recently, in two occasions in September 2017 a local resident of the small locality Media Luna (lat. −18.9533, lon. - 65.1384) in the Rio Chico valley (Fig. 3), has reported specimens to the health centers of La Palma (about 50 km by road, south of Estancia Mataral), after discovering them in its peridomicile. Then, in April 2020 we collected another specimen in a domicile (lat. −18.8255, long. −65.1194) in Chuqui Chuqui, a small locality from the Rio Chico valley at about 15 km north of La Palma. In that instance, it was merely a unique specimen that had entered the house, lured by indoor lighting. In the same year, shortly afterward, another specimen was brought to the health center by an inhabitant of Chuqui Chuqui. The three collecting points of Bolivia are in close proximity. The valleys of Rio Chico and Rio Novillero converge at nearly the same point along the Rio Grande, with one valley seamlessly continuing into the other in a south-to-north direction. The straight-line distances between key locations are as follows: Mataral Estancia to Chuqui Chuqui is about 25 km, and Chuqui Chuqui to Media Luna (La Palma) is about 15 km. Considering the two nearest occurrences between Bolivia and Argentina, it becomes evident that the distance spans approximately 950 km, indicating a substantial gap between them (Fig. 1).

**Fig. 3.**
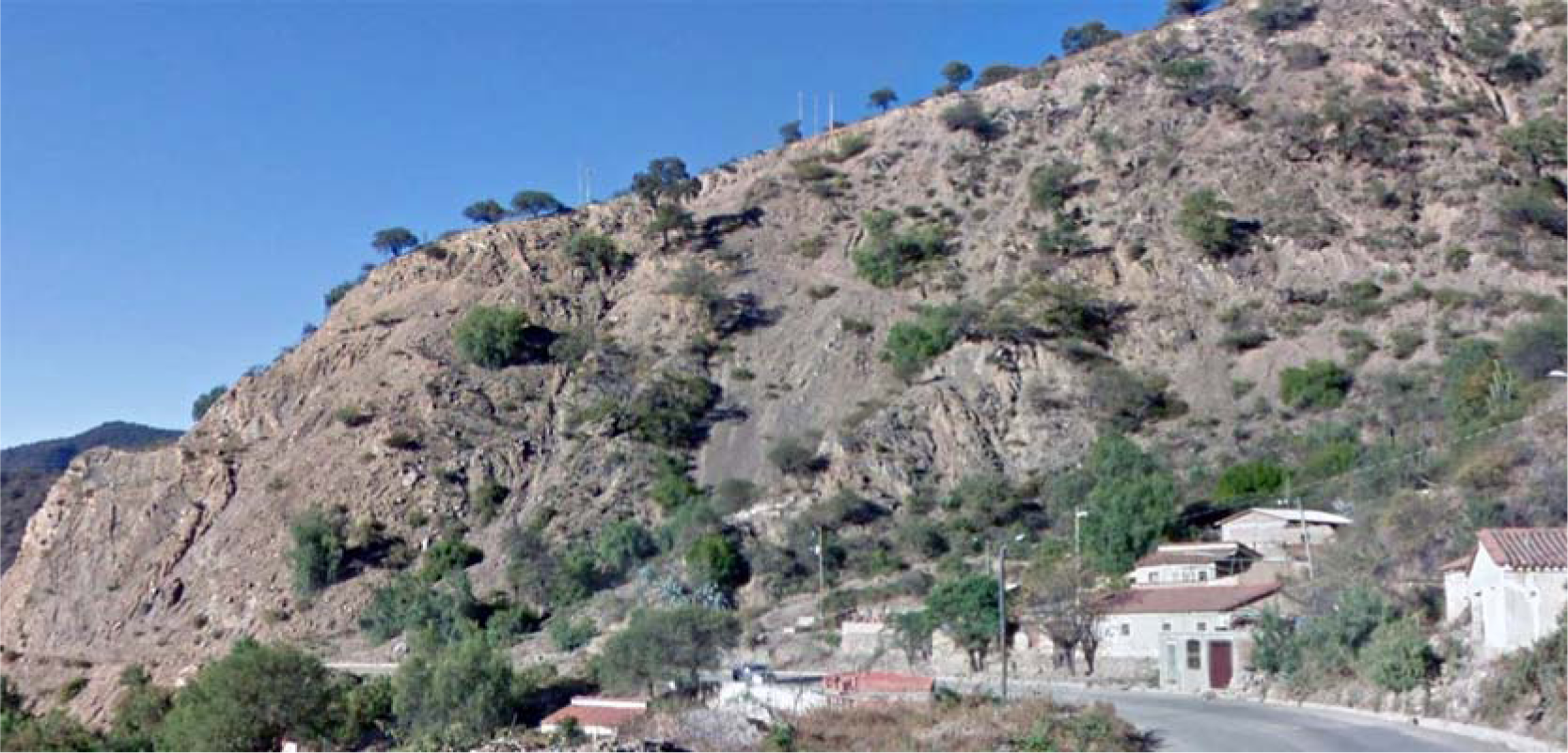
Collecting area in Media Luna locality (Bolivia), showing the general environment and the house where *T. breyeri* was collected.

The three discovery points in Bolivia lies within the WWF “Bolivian Monte Dry Forest” ecoregion [17] and are located in the Köppen-Geiger class “BSh” (arid, steppe, hot) characterized by a pronounced dry season which can last several months. This climate class is also the one where most Argentinean locations are situated. The ecoregion is in the dry mountain valleys of the Andes in southern Bolivia. There, the mean monthly temperatures range from 13.5 C in July to 18.8°C in November. Yearly total precipitation is about 580 mm. Average monthly precipitation is below 10 mm in May–August, and rises to 137.4 mm in January.

### Identification

#### Morphological identification

Identification was conducted using unambiguous identification keys and descriptions [2, 3]. *Triatoma breyeri* closely resembles *T. eratyrusiformis* but can be distinguished by the absence of long hairs on the appendages, rounded posterior angles of the thorax, and uniform coloration along the lateral edges of the abdomen [27]. A general view of one of the Bolivian specimens from Estancia-Mataral is given in Fig. 4 and S1, S2, S3 and S4 Figure.

**Fig. 4.**
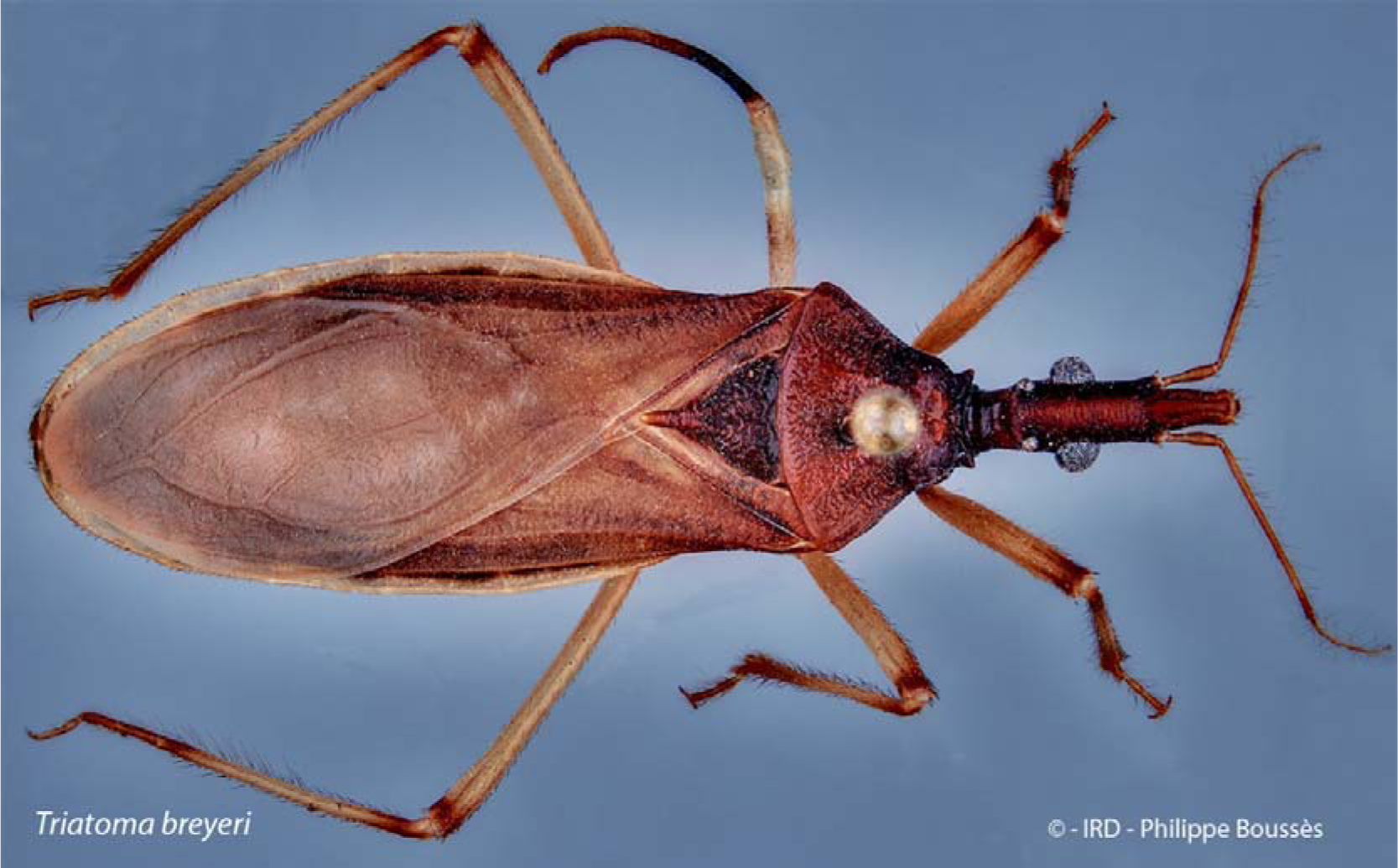
General dorsal view of a *T. breyeri* specimen collected in Estancia-Mataral, Bolivia (voucher “Isolate A”).

#### Molecular identification

Two specimens of *T. breyeri* were used. The first one (“isolate A”) from Estancia-Mataral and the second one (“isolate B”) from Chuqui Chuqui. The two voucher specimens for the samples sequenced in this study were deposited in the collection of the Entomological Laboratory (LEMUMSS) of the Universidad Mayor de San Simón at Cochabamba, Bolivia, under references LMUR-2022-01 and FLX-2019-01 respectively.

Results for sequences of cytochrome b (*CytB*), cytochrome c oxidase subunit I (*COI*) and 16S ribosomal RNA (*16S*) genes were as follow:

##### CytB

Isolates A and B presented sequences of 319 bp, only separated by one mutation at position 97. These sequences were deposited in GenBank with numbers OR664766-7. The BLAST search using both A and B sequences as query and limiting the hits to a qc > 95% and an id > 84% returned 31 hits resulting in 21 unique sequences. The most closely related sequences were the isolate 56, a *T. breyeri* specimen from Mataral, Cochabamba, Bolivia, accession number KC249242.1 [28], presenting identities of 100% and 99.69% with A and B respectively, then KC236980.1 [29] and JN102361.1 [7], both bugs labeled *T. breyeri* and isolated from Patquia Viejo, La Rioja, Argentina and presenting identities of 95.92% and 96.24% with our isolates A and B respectively. In the ML tree, these sequences are clustered into a significant monophyletic group (bootstrap value of 100 (S5 Figure), whose most closely related sister clade was *Triatoma maculata*.

##### COI

A and B isolate sequences of 355 bp (GenBank numbers OR668702-3) differed by three mutations at positions 108, 282 and 333. The BLAST search using A and B as query only limited by a qc > 95% returned 15 hits resulting in 14 unique sequences. Isolate A sequence is identical to the *T. breyeri* reference KC249321.1 [28], from the isolate 56 of Mataral, Cochabamba, Bolivia while and isolate B presented an identity 99.15% with that GenBank reference. In the ML tree, these three sequences are closely related and clustered with *Triatoma eratyrusiformis* haplotypes in a significant monophyletic group (bootstrap value of 99) (S6 Figure), the latter being a sister clade to the *Mepraia spinolai* cluster.

##### 16S

Sequences of 328 bp from A and B isolates were identical (GenBank numbers OR668706-7). The BLAST search using this unique haplotype as query and limited by qc > 95 and id > 91 returned 41 hits resulting in 25 unique sequences. This haplotype showed 99.70% identity with *the T. breyeri* reference KC248988.1, reported from the isolate 56 of Mataral, Cochabamba, Bolivia [28]. In the ML tree (S7 Figure.) our isolates and the *T. breyeri* reference were clustered into a significant group (bootstrap value of 90) closely related to a *T. eratyrusiformis* reference, all being a sister clade of the *Mepraia spinolai* group.

### MaxEnt model of habitat suitability

The MaxEnt model was constructed using the *kuenm* R-package [30]. The *kuenm* process selected 46 statistically significant MaxEnt models. The top-performing tenth of the models were constructed employing the bioclimatic variables BIO6 (minimum temperature of the coldest month), BIO13 (precipitation of the wettest month), and BIO17 (precipitation of the driest quarter), with minimal variations attributed solely to differences in regularization parameters or feature classes. The foremost model was characterized by a regularization multiplier set at 0.6; and feature class ‘l’. This model was ultimately selected as the definitive model. The computed mean AUC value of 0.93 represent an excellent performance and prediction of the model [31] (general MaxEnt outputs are in S1 Table). The representation of the habitat suitability is given in Fig. 5. The Complementary Log-Log (cloglog) output format of MaxEnt was used. It gives an estimate between 0 and 1 of probability of presence of the species [32]. The median prediction value from all 12 model replicates (from bootstrapping) is represented in Figure 5. Unlike the mean, the median has a lower sensitivity to extreme values, which could be prevalent across replicates. Nevertheless, for reference, the mean, minimum and maximum predictions are available within the S8 Figure, S9 Figure and S10 Figure respectively. To enhance the clarity of the *T. breyeri* habitat suitability maps, the suitability threshold chosen for visualization was the value of the mean maximum training sensitivity plus specificity (Max TSS) Cloglog. Consequently, areas with values below this threshold (0.6144 in this study) are depicted in grey in Figure 5.

**Fig. 5.**
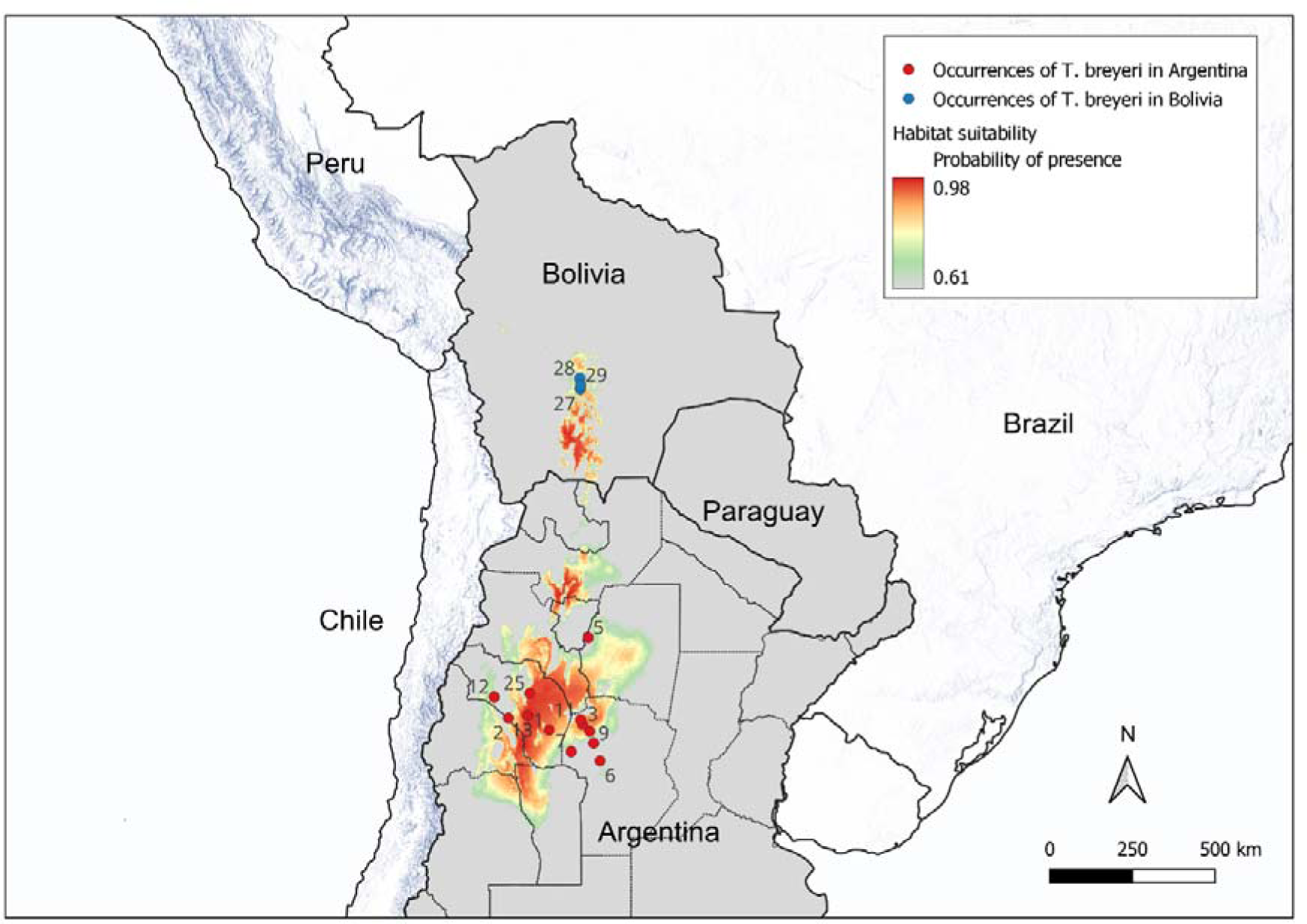
MaxEnt representation of habitat suitability for *T. breyeri.* Median predictions. MaxEnt output is the Complementary Log-Log (cloglog) format which is a probability of presence. Median values of 12 model replicates (boostrap) are computed. The mean maximum sensitivity plus specificity (MaxTSS) Cloglog threshold was used to depict in grey values below it (0.63 in the study). Numbers in the map refer to Id number of Table 1.

Habitat suitability for *T. breyeri* encompasses approximately the ecoregions of “Dry Chaco” and “High Monte” in Argentina and, not surprisingly, the southern part of the “Bolivian Montane Dry Forest” where it was collected in Bolivia. In this region, the prevailing Köppen-Geiger climate classes are BSh (arid, steppe, hot) and BSk (arid, steppe, cold). It is worth noting that these specific climate classes correspond to the majority of *T. breyeri* occurrences found in Argentina.

**Table 1.** Geographical characteristics of *T. breyeri* occurrences in Argentina and Bolivia. ID numbers correspond to those in the MaxEnt representation of habitat suitability in Fig. 5.

| ID | Country | Locality | Provincia (Argentina) o Departamento (Bolivia) | Latitude | Longitude | Collecting Date | Source | Köppen-Geiger climate classification |
| --- | --- | --- | --- | --- | --- | --- | --- | --- |
| 1 | Argentina | Iliar | La Rioja | -30.67017 | -66.20836 | 1940 | GBIF | BSh |
| 2 | Argentina | Salinas de Bustos | La Rioja | -30.26333 | -67.60611 | 2008 | GBIF | BWk |
| 3 | Argentina | Guanaco Muerto | Cordoba | -30.47905 | -65.06132 | 1977 | GBIF | BSh |
| 5 | Argentina | Termas de Rio Hondo | Santiago del Estero | -27.49855 | -64.86053 | 1977 | GBIF | Cwa |
| 6 | Argentina | La Serranita | Cordoba | -31.73501 | -64.45888 | 1978 | GBIF | Cwa |
| 7 | Argentina | Chancani | Cordoba | -31.41621 | -65.45018 | 1989 | GBIF | BSh |
| 9 | Argentina | Cordoba | Cordoba | -31.12600 | -64.67800 |  | GBIF | Cwb |
| 10 | Argentina | Cruz del Eje | Cordoba | -30.72185 | -64.80864 |  | GBIF | BSh |
| 11 | Argentina | Los Leones | Cordoba | -30.31800 | -65.12322 |  | GBIF | BSh |
| 12 | Argentina | Pagancillo | La Rioja | -29.54111 | -68.09690 | 1970 | GBIF | BWk |
| 13 | Argentina | Patquia Viejo | La Rioja | -30.17342 | -66.95070 | 2009 | GBIF | BWh |
| 25 | Argentina | La Rioja | La Rioja | -29.40733 | -66.86652 |  | GBIF | BWh |
| 27 | Bolivia | Media Luna | Sucre | -18.95336 | -65.13846 | 2017 | This study | BSh |
| 28 | Bolivia | Estancia Mataral | Cochabamba | -18.59638 | -65.14842 | 2010 | This study | BSh |
| 29 | Bolivia | Chuqui Chuqui | Sucre | -18.82558 | -65.11938 | 2020 | This study | BSh |

It is worth noting that the three bioclimatic variables, BIO6, BIO13, and BIO17, do not have equal contributions to the model construction. BIO13 exhibits a relatively low contribution value of 0.83, whereas BIO17 and BIO6 contribute significantly with values of 41.5 and 57.6, respectively. This discrepancy is further evident in their permutation importance, with values of 1.47, 41.9, and 56.6 for BIO13, BIO17, and BIO6, respectively (S1 Table).

## Discussion

### Species molecular identification and clustering analysis

The three gene fragments used here unambiguously characterize isolates A and B as *T. breyeri*. In the case of *16S* and *COI*, it is unsurprising that isolate A and B clustered together with “isolate 56 Nb. seq. 1”, a specimen also originating from Mataral, Bolivia [8]. However, it’s worth noting that when analyzing the *CytB* tree, which included two insects from Argentina, these Argentinean insects unequivocally clustered with the Bolivian specimens.

The species is very close to *T. eratyrusiformis* and to a lesser extent to *M. spinolai*, as mentioned by morphological taxonomists [3] a correlation further supported by the *COI* and *16S* trees of this study. Unfortunately, there was no *CytB* sequence for these last two species which would have made it possible to confirm this trend. Nevertheless, the data presented here do not allow definitive conclusions to be drawn on the exact phylogenetic position of *T. breyeri*.

### Ecology

The consistent capture of *T. breyeri* across multiple sites since 2010 in the region of Estancia-Mataral – La Palma provides substantial evidence that the species has established a robust presence in the area. At the country level, the reporting of Triatomines at healthcare facilities is a common practice in Bolivia, with community reports playing a vital role in optimizing the scheduling of insecticide sprayings by the National Chagas Program. Despite this extensive community involvement and our collections, there have been no reports of *T. breyeri* from other regions of Bolivia. Currently it appears that the presence of this species is limited to the studied region comprised in the small valleys of Rio Chico and to a lesser extend of Rio Novillero.

The persistence of the species in this limited region has occurred despite the insecticide interventions carried out to control *T. infestans.* Notably, the insecticide applications primarily target domiciles and peri-domiciles, while *T. breyeri* is known to utilize sylvatic breeding sites, although it is also found in peri-domiciliary environments. The low prevalence of the species may also be attributed to other bio-ecological factors. In the region, *T breyeri* may experience interspecific competition (although probably reduced) from *T. guayazayana*, and *T. sordida*, which are two other sylvatic/peridomestic species. In the domiciles (which are not its usual biotopes), it will encounter *T. infestans,* a very competitive species that is the most abundant in the region. Furthermore, *T. breyeri* exhibits a replacement rate of only one-third of that of *T. infestans* [25] which in not in favor of a rapid population growth.

The potential geographic distribution of the species, as indicated by the MaxEnt model, suggests that Bolivia may represent a habitat enclave within the species’ extreme geographical range. The best MaxEnt model predicting habitat suitability was constructed with a limited number of occurrences in Argentina. While this number is low, the number of occurrences was sufficient for accurate results [33]. Although the model appears accurate, it should be interpreted with caution, yet it provides valuable insights. The model points out that both temperature and precipitation variables play a role in the species habitat suitability, as previously mentioned [34]. In particular, within the selected set of climatic variables, the minimum temperature of the coldest month (BIO6) and the precipitations of the driest quarter (BIO17) play a predominant role. The model highlights the suitability of the “Bolivian Montane Dry Forest” ecoregion, characterized by Köppen-Geiger climate classes BSh and BSk. These classes are akin to the climate conditions found in the nearby “High Monte” ecoregion of Argentina, where *T. breyeri* is known to exist. Despite being categorized as “arid and steppe,” the Bolivian region where *T. breyeri* is present exhibits a cooler climate, which might serve as a potential limiting factor for its distribution in the area. Considering all these bio-ecological factors, the species appears to persist in very low densities in Bolivia.

As of today, the species has not expanded its range to encompass the entire ‘Bolivian Montane Dry Forest’ ecoregion. This limitation is likely due to the ecoregion serving as an ecological distribution boundary, as well as the species’ recent introduction and relatively slow development rate. However, there are occurrences in various distinct locations within its limited presence in Bolivia, possibly because the species exhibits a good flight dispersal capacity [25], allowing it to gradually colonize new sites.

The infrequent occurrence of *T. breyeri* in Bolivia and the substantial distance separating its known presence from its focal points of presence in Argentina strongly suggest that the species was likely introduced into the region relatively recently, and probably as a result of accidental introduction. Passive transport of the species by a vertebrate host (including humans) is a reasonable hypothesis [35]. As the species exhibits a high longevity and resist fasting [25], it can therefore withstand travel inside human belongings.

*Triatoma breyeri* remains a species of considerable mystery, with its presence in Bolivia marked by limited knowledge and few collection points. Further research and monitoring efforts are essential to gain a more comprehensive understanding of this species, its behavior, and its potential implications for the region.

## Material and methods

### Molecular identification

Two samples (isolate A from Estancia-Mataral and isolate B from Chuqui Chuqui) were processed to genomic DNA extraction, from one hind leg, using the Qiagen DNeasy Blood & Tissue kit according to the manufacturer instructions except that insect’s leg have been manually homogenized in 180µl of PBS 1X with a pestle and incubated 3 hours with 20µl of proteinase kinase and 200µl of AL buffer, vortexed multiple times during incubation. Amplification of the *16S* ribosomal RNA, cytochrome c oxydase subunit 1 (*COI*) and cytochrome b (*CytB*) genes from genomic DNAs was carried out by simplex PCR with respective primers 16S_F (5’-GGG CTG GAA TGA AAG GTT GG-3′) and 16S_R (5′-GCT CCT GCA CCC ACA ACT TA-3′), COI_F (5’-ACG TTG ACA CAC GAG CAT AC-3’) and COI_R (5’-AAG TGT TGT GGG AAG AAT GTC A-3’), CytB_F (5’-TCT GCG ATC CCT TAC TTA GGA-3’) and CytB_R (5’-TCA GGT TGA ATG TGT ACT GGG G-3’) [7, 8, 36].

The PCR reaction mix using the Qiagen Taq DNA Polymerase kit, was prepared as follow: 1X buffer (containing 1.5mM of MgCl2), 0.5mM of MgCl2, 0.2mM of dNTP, 0.2µM of both forward and reverse primers and 1U of taq polymerase. The reaction mix was performed with 2μL of sample DNA for a final volume of 25µl. PCR reactions of three loci were performed using the same following cycling conditions: an initial denaturation at 94°C for 2 min, 10 cycles at 94°C for 30 sec, 65°C [-1°C/cycle] for 30 sec and 72°C for 30 sec, 20 cycles at 94°C for 30 sec, 55°C for 30 sec and 72°C for 30 sec, and a final extension at 72°C for 5 min. For sample A and *CytB* gene, a specific program was done with following cycling conditions: an initial denaturation at 94°C for 2 min, 30 cycles at 94°C for 30 sec, 55°C for 30 sec and 72°C for 30 sec, and a final extension at 72°C for 5 min.

The expected size of the PCR products (including the primers) of 369, 397 and 372 base pairs (bp) for *16S* rRNA, *COI* and *CytB* genes respectively, were confirmed by electrophoresis on a 1.5% agarose gel stained with GelRed and visualized by UV transillumination.

Sequencing was performed by Eurofins Genomics sequencing services. Sequences of all amplicons (*CytB*, *COI* and *16S*) were manually corrected from Sanger’ chromatograms and primers were removed. Each sequence was compared by BLAST searching against the database of Triatominae (taxid:70999) on the NCBI site. The hits returned were limited by a minimum query cover (qc) > 95% to remove sequences that were too short and by a minimum identity (id) depending on the quantity of sequences returned, in order to build a readable tree. The best model of nucleotide substitution according to the lowest BIC (Bayesian Information Criterion) score was estimated and a maximum likelihood tree (ML tree) was constructed with this best model. A phylogeny statistical test based on bootstrap [37] of 100 replicates was applied for each locus. Sequence alignments, best model determinations and ML trees were performed with MEGA7 [38].

### Habitat suitability

Habitat suitability analysis for *T. breyeri* was conducted through the application of an ecological niche modeling approach employing the MaxEnt algorithm (ver. 3.4.4) [39, 40] and following established recommendations for good practice [41–43]. The analysis involved the following sequential steps:

#### Occurrence data

Occurrence data for *T. breyeri* in Argentina were obtained from the Global Biodiversity Information Facility (GBIF: https://gbif.org). The extracted dataset comprised of 181 occurrences (GBIF.org (28 August 2023) GBIF Occurrence Download https://doi.org/10.15468/dl.syhe92). The data underwent cleaning to remove repeated observations, ensure sufficient decimal precision in latitude and longitude coordinates, verify their presence within the expected region (Argentina), and identify potential outliers. This process resulted in a dataset of 12 occurrences (Table 1) sourced from the DataTri database [44].

Spatial autocorrelation within the dataset of occurrences indicates the lack of independence among geographically close observations, which could potentially influence model outcomes. To assess the spatial autocorrelation among occurrences, the *variogmultiv* function from the R-package *adespatial* was employed, considering the five selected bioclimatic variables (see next paragraph). The analysis indicated the absence of significant autocorrelation, eliminating the necessity to thin the occurrence dataset by removing geographically close points. Consequently, all 12 occurrences were retained. The direct line distances between these 12 Argentinean points ranged from 19 to 471 km.

#### Environmental predictors

The initial set of the 19 bioclimatic variables (BIO1 to BIO19) from the WorldClim ver. 2.1 database [45] at a resolution of 2.5 arc-minute (approximately 5 km) represented the various candidate predictors that are potentially relevant for the geographic distribution of *T. breyeri* (data available at: https://www.worldclim.org/data/worldclim21.html). This dataset is representative of annual and seasonal means variations of temperatures and precipitations averaged over a 30-year period (1970-2000).

#### Selection of relevant environmental predictors

Because there exist some artifacts in the WorldClim variables BIO8, BIO9, BIO18 and BIO19 [46, 47], they were excluded from the pool of selected variables to build the models. When constructing models with collinear variables, there is an elevated risk of overfitting and over-parametrization [48]. The reduction of relevant variables was therefore achieved through statistical analysis, with a focus on retaining those variables that held biological significance for Triatomines. The 15 remaining WorldClim variables were submitted to a VIF (Variance Inflation Factor) analysis using the R-package *usdm* [49]. In the package, the two functions *vifcor* and *vifstep* were used with recommended thresholds of 0.7 [50] and 5 [51, 52] respectively. The function *vifcor*, first find a pair of variables which has the maximum linear correlation (greater than the threshold), and exclude one of them which has greater VIF. The procedure is repeated until no variable with a high correlation coefficient greater than the threshold with other variables remains. The function *vifstep* calculate VIF for all variables, exclude one with highest VIF (greater than threshold), repeat the procedure until no variables with VIF greater than the threshold remains. The functions were applied to the pixel values of all 15 bioclimatic raster layers within the area defined by a convex hull around the 12 occurrence records in Argentina. The two VIF analysis consistently selected the variables BIO2, BIO4, BIO6, BIO13 and BIO17 (S2 Table).

#### Model calibration

Maxent exhibits proficiency in managing small and sparsely populated datasets, albeit requiring meticulous parameter refinement [53–56]. This fine-tuning process entails the adjustment of critical settings [42, 57], encompassing feature selection, regularization multipliers, convergence criteria, and the selection of pertinent predictor variables, all directed toward the optimization of the model’s performance. To carry out such model calibrations, the R-package *kuenm* [30] was used with the following settings: (1) The function *kuenm_occsplit* was used to generate the data sets selecting at random 80% of the occurrences for the training data set; (2) In the function *kuenm_cal*, the regularization multipliers to be evaluated range from 0.1 to 1 with an increment of 0.1 and from 1.5 to 6 with an increment of 0.5 and values of 8 and 10 *(i.e.,* a total of 22 regularization parameters). It is worth noting that the optimal value may lie within the extremes of this range [58, 59]. In that function, the feature class to be evaluated were linear (*l*), the combination linear-quadratic (*lq*), hinge (*h*), and the combination *lqh* (*i.e.,* a total of 4 feature class combinations); (3) From the five selected climatic variables, multiple variable sets were generated using the *kuenm_varcomb* function, each set containing a minimum of 3 variables. Consequently, from the initial selection of BIO2, BIO4, BIO6, BIO13 and BIO17, a total of 16 distinct sets were created.

In all, 1408 candidate models were evaluated (combinations of 22 regularization multiplier settings, 4 feature class combinations, and 16 distinct sets of environmental variables). Model performance was evaluated on significance (partial *ROC*, with 500 iterations and 20% of data for bootstrapping), omission rates (*OR*, with parameter *E*=5% [60]), and the Akaike information criterion corrected for small sample sizes (*AICc*).

#### Final models

Final models were created using the *kuenm_mod* function of the <u>kuenm</u> R-package, with parameters settings of 12 replicates using recommended bootstrapping [61]. A jackknife process was executed to evaluate the relative importance of single explanatory variables included in the models.

## Author contributions

### Conceptualization

Frédéric Lardeux

### Data curation

Frédéric Lardeux, Christian Barnabé, Lineth Garcia

### Formal Analysis

Frédéric Lardeux, Christian Barnabé

### Funding acquisition

Frédéric Lardeux

### Investigation

Frédéric Lardeux, Alberto Llanos, Roberto Rodriguez, Christian Barnabé, Luc Abate, Philippe Boussès, Rosenka Tejerina-Lardeux, Lineth Garcia

### Methodology

Frédéric Lardeux, Christian Barnabé, Luc Abate

### Resources

Frédéric Lardeux, Luc Abate, Alberto Llanos, Philippe Boussès, Rosenka Tejerina-Lardeux, Roberto Rodriguez, Lineth Garcia

### Validation

Frédéric Lardeux, Luc Abate, Alberto Llanos, Philippe Boussès, Rosenka Tejerina-Lardeux, Roberto Rodriguez, Lineth Garcia

### Supervision

Lineth Garcia, Frédéric Lardeux

### Writing – original draft

Frédéric Lardeux, Christian Barnabé, Luc Abate, Philippe Boussès

### Writing – review & editing

Frédéric Lardeux, Christian Barnabé, Luc Abate, Alberto Llanos, Philippe Boussès, Rosenka Tejerina-Lardeux, Roberto Rodriguez, Lineth Garcia

## Supporting information

**S1 Figure.**
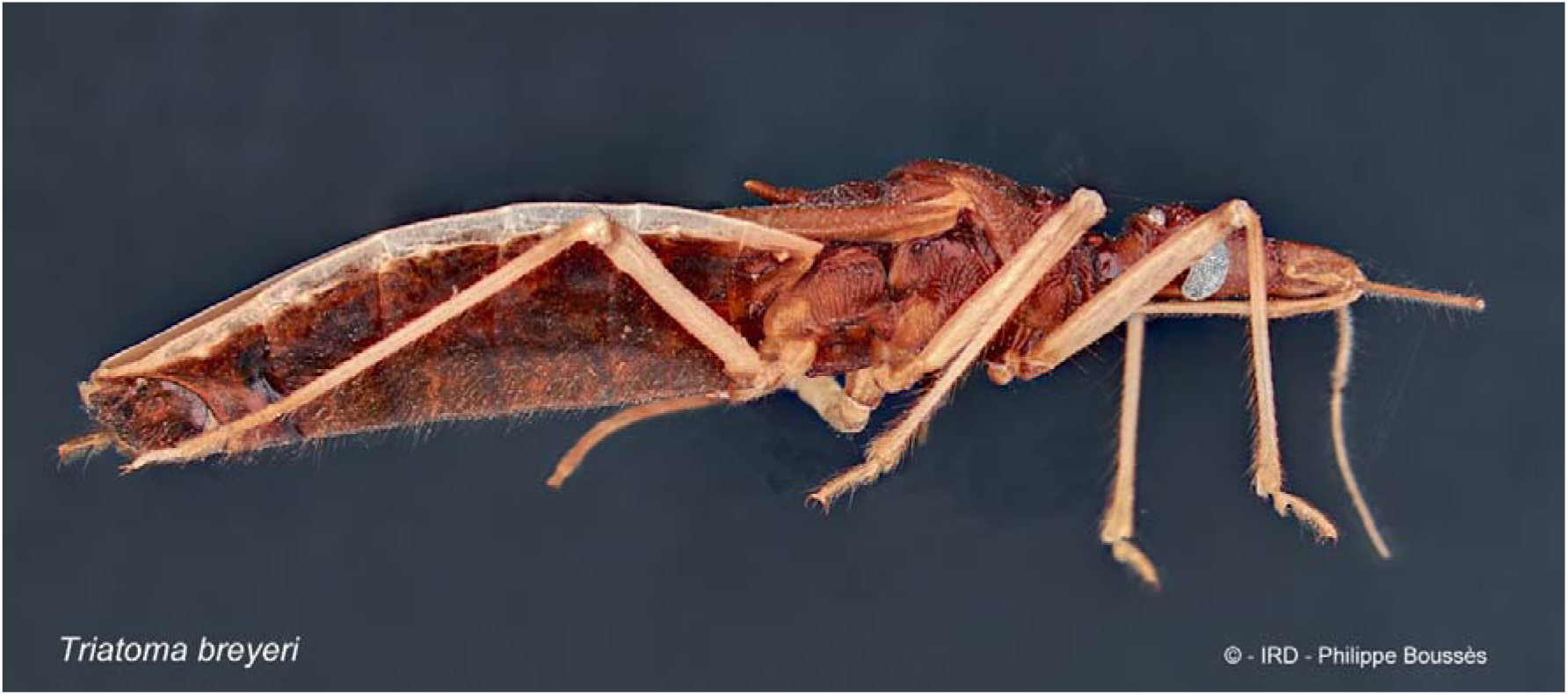
Lateral view of a specimen of *Triatoma breyeri* from Estancia-Mataral, Bolivia. The specimen depicted is the one used for molecular identification and is named isolate A in the present study.

**S2 Figure.**
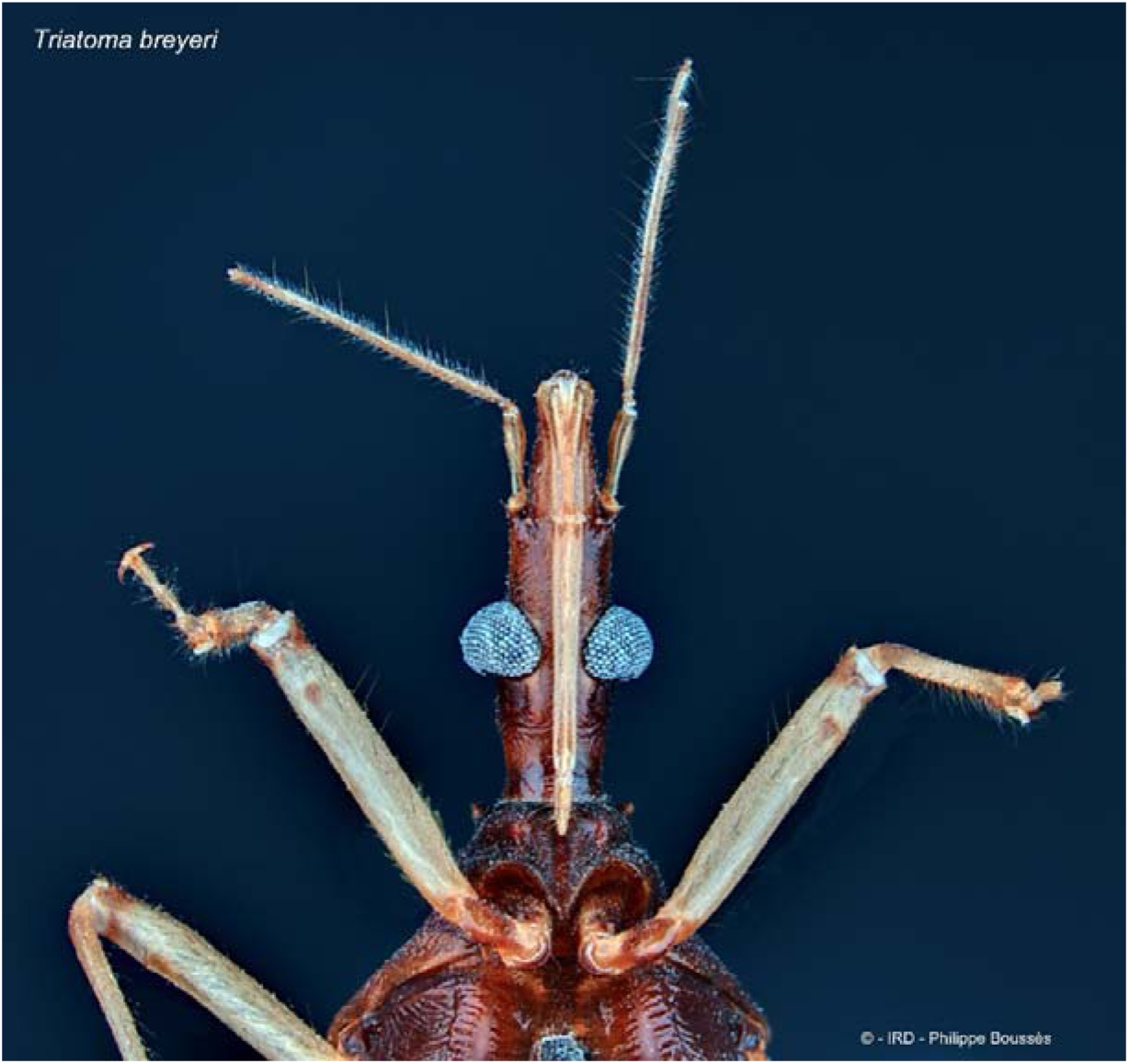
Dorsal view of the head of a specimen of *Triatoma breyeri* from Estancia-Mataral, Bolivia. The specimen depicted is the one used for molecular identification and is named isolate A in the present study.

**S3 Figure.**
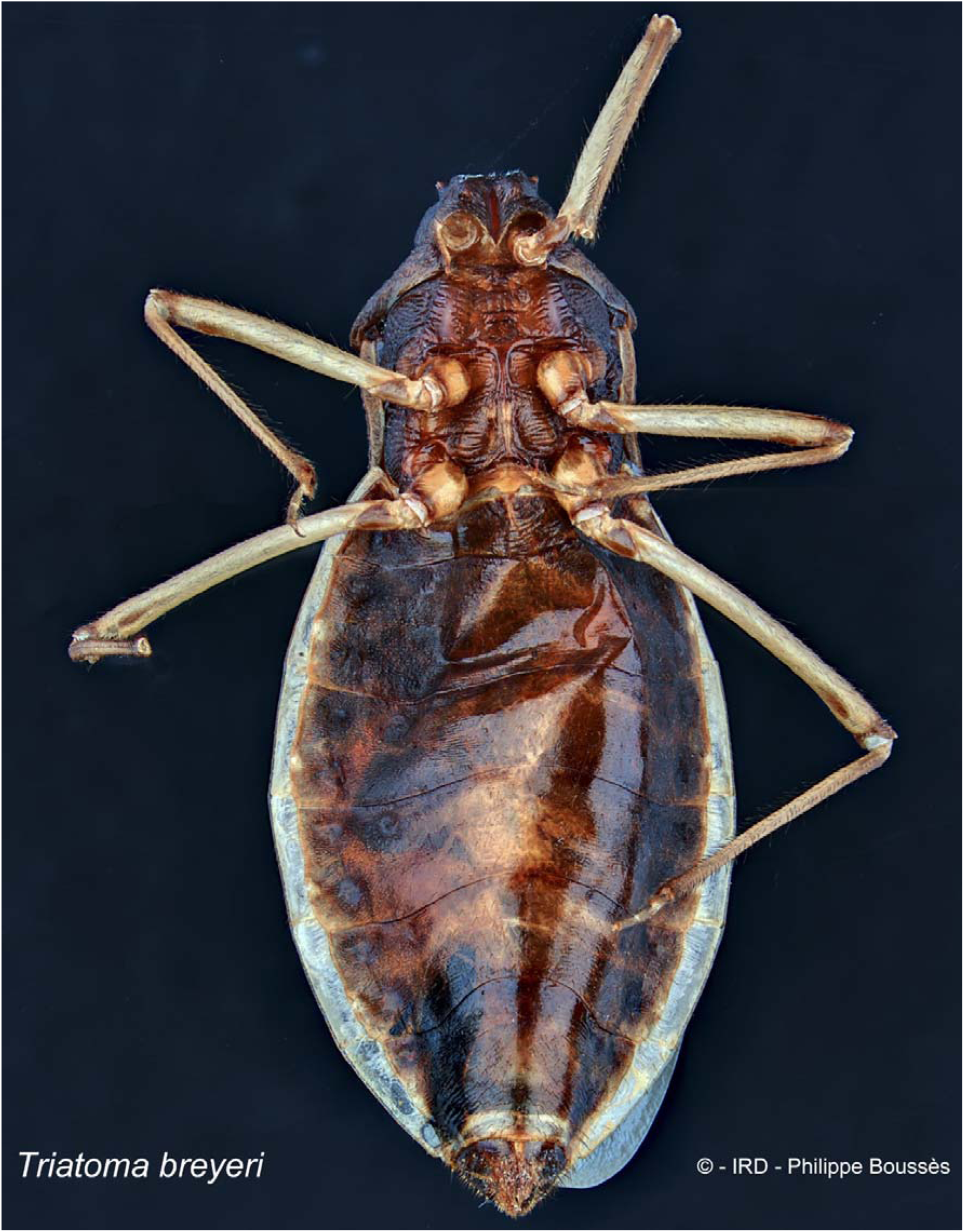
Ventral view of a specimen of *Triatoma breyeri* from Chuqui-Chuqui, Bolivia. The specimen depicted is the one used for molecular identification and is named isolate B in the present study. Head is missing.

**S4 Figure.**
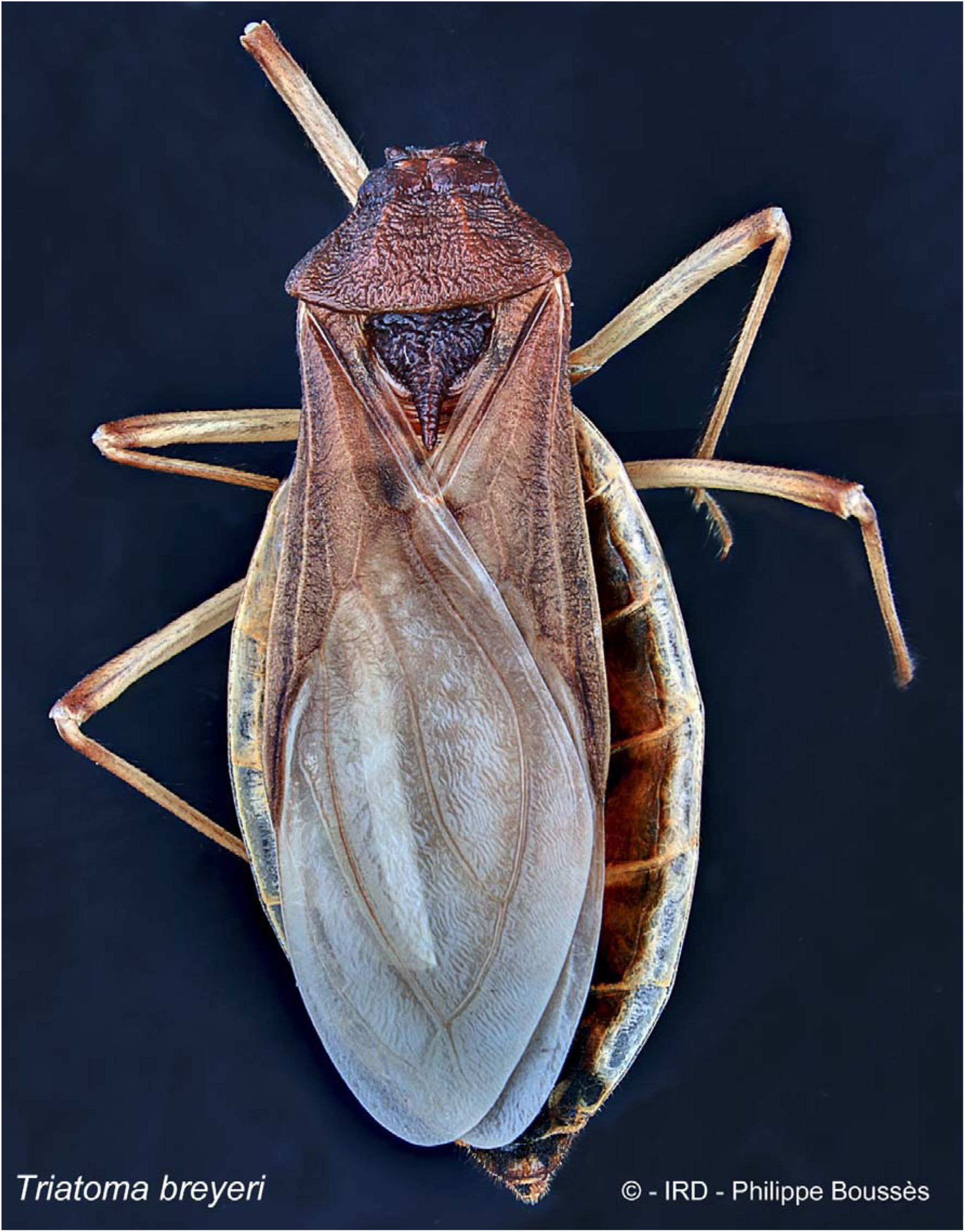
Dorsal view of a specimen of *Triatoma breyeri* from Chuqui-Chuqui, Bolivia. The specimen depicted is the one used for molecular identification and is named isolate B in the present study. Head is missing

**S5 Figure.**
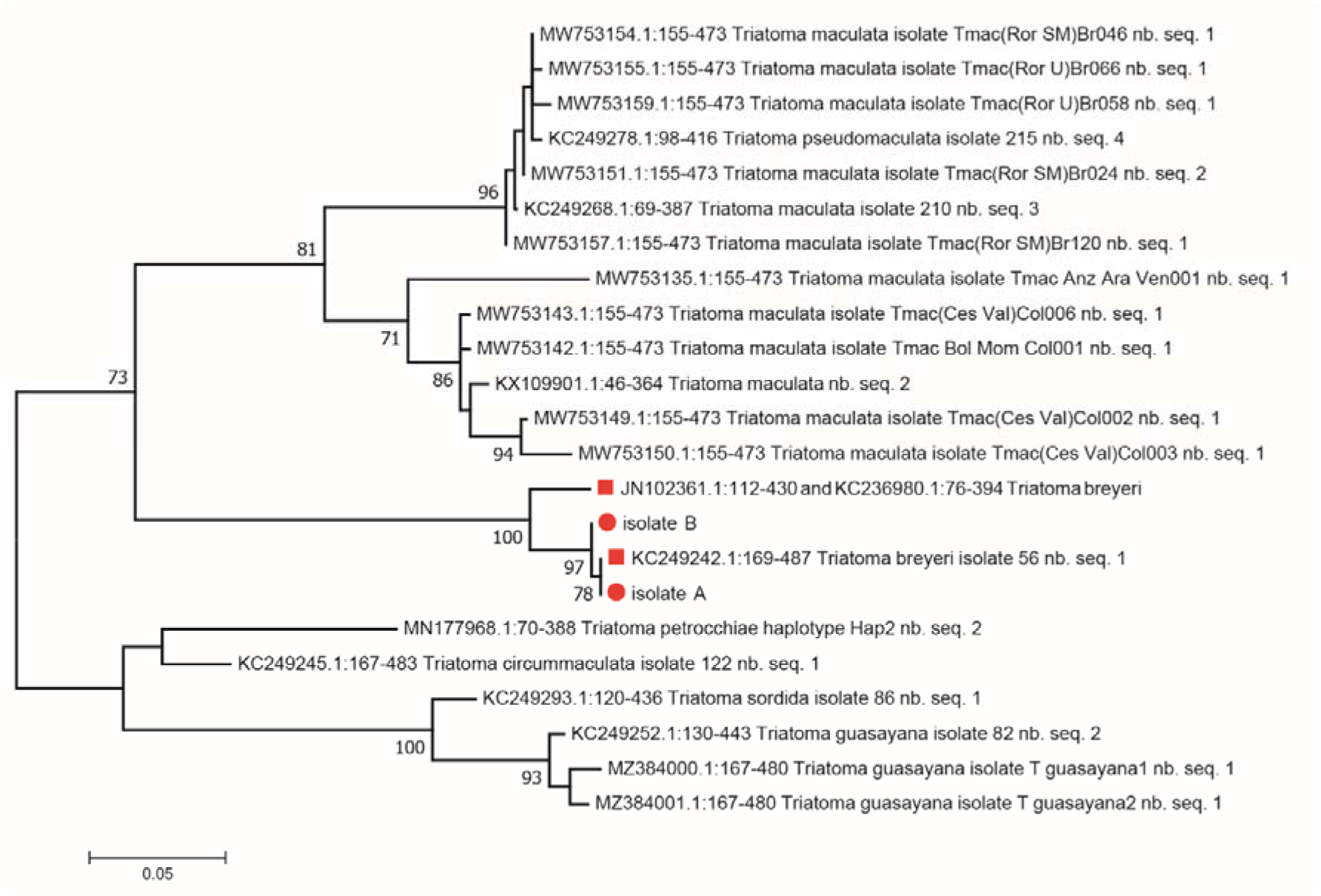
Maximum likelihood (ML) tree of 23 different *CytB* haplotypes constructed from an alignment of 319 nucleotide positions. The numbers at the nodes correspond to the bootstrap values > 70%. The best multiple substitution model was the Tamura-Nei model +G (Gamma distribution) + I (invariants). Isolates A and B are visualized as red circles while *T. breyeri* reference sequences are represented as red squares. “nb. seq.” is the amount of the particular haplotype in GenBank.

**S6 Figure.**
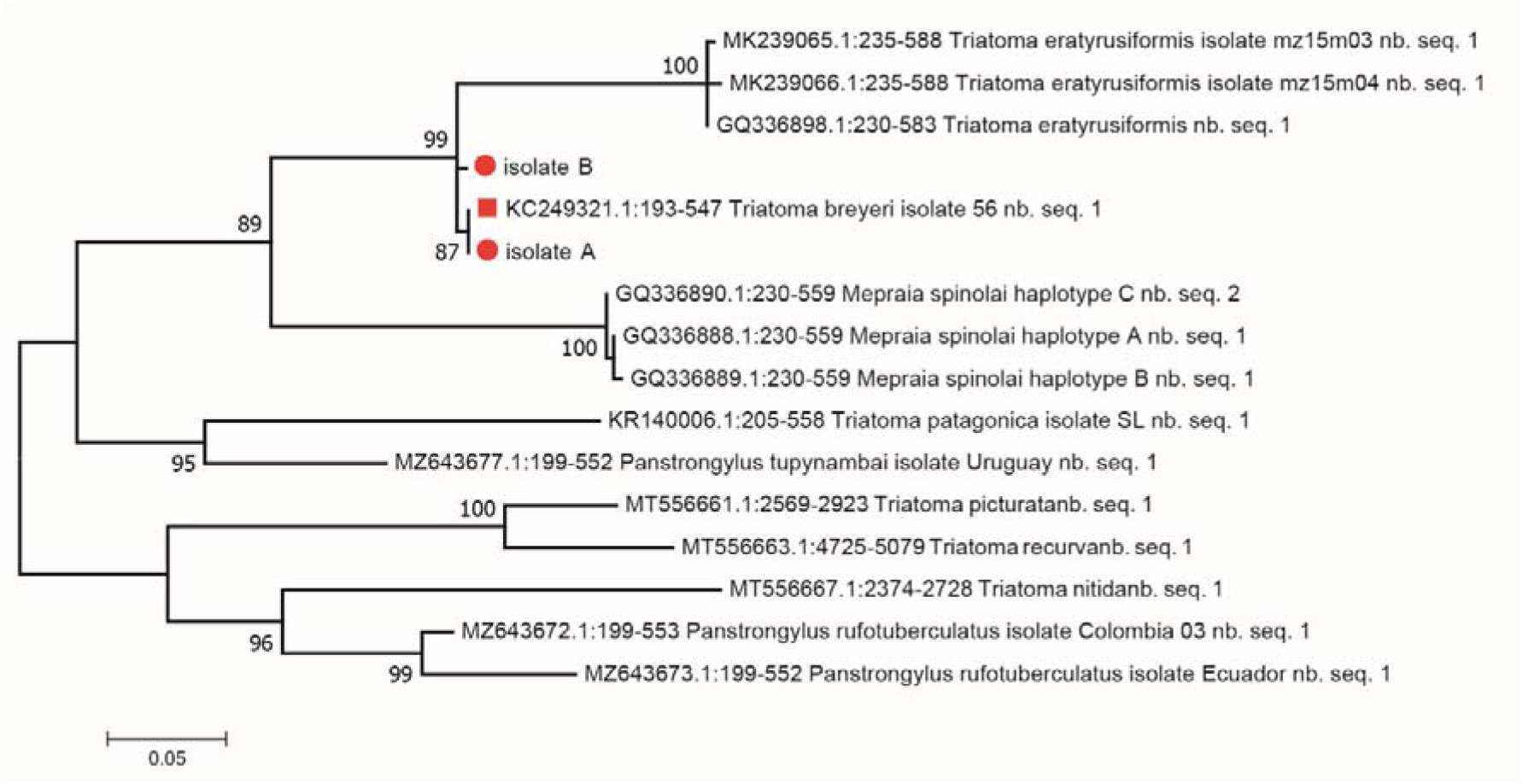
Maximum likelihood (ML) tree of 16 different *COI* haplotypes constructed from an alignment of 354 nucleotide positions. The numbers at the nodes correspond to the bootstrap values > 70%. The best multiple substitution model was the GTR model +G (Gamma distribution) + I (invariants). Isolates A and B are visualized as red circles while the only *T. breyeri* reference sequence is represented as a red square. “nb. seq.” is the amount of the particular haplotype in GenBank.

**S7 Figure.**
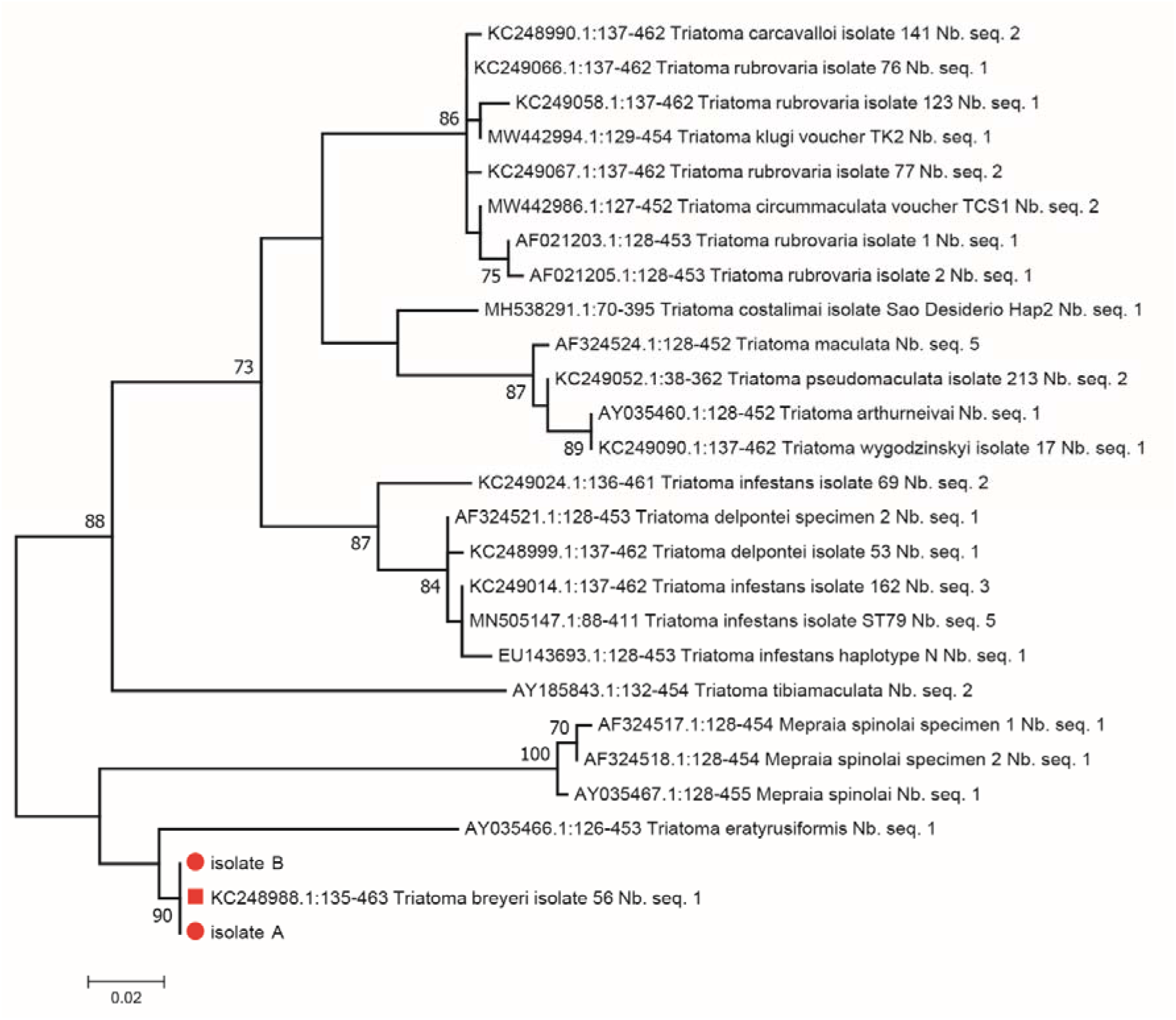
Maximum likelihood (ML) tree of 27 different *16S* sequences constructed from an alignment of 326 nucleotide positions. The numbers at the nodes correspond to the bootstrap values > 70%. The best multiple substitution model used was the Hasegawa-Kishino-Yano mode (HKY) model +G (Gamma distribution). Isolates A and B are visualized as red circles while the only *T. breyeri* reference sequence is represented as a red square. “nb. seq.” is the amount of the particular haplotype in GenBank

**S8 Figure.**
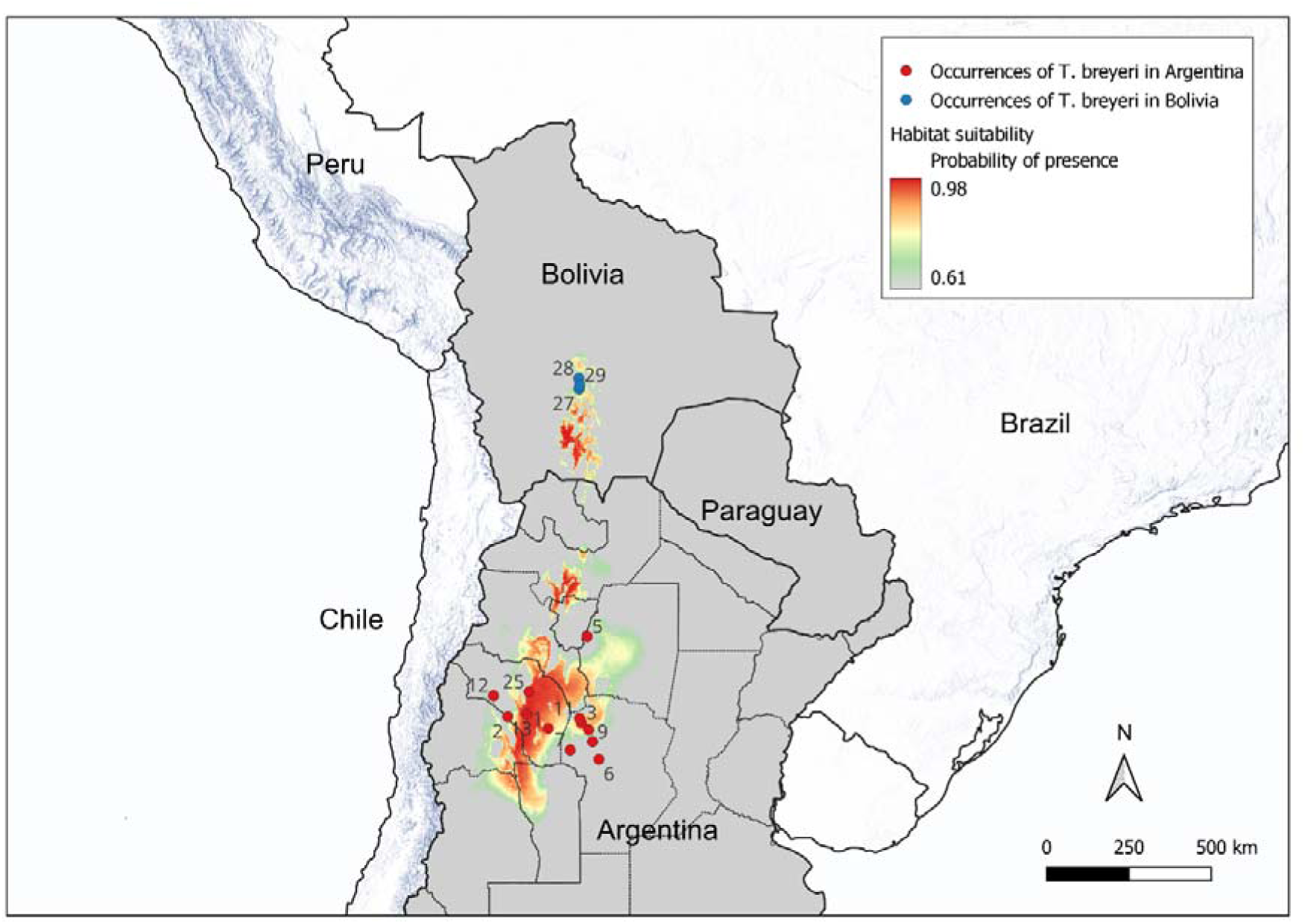
MaxEnt representation of habitat suitability for *T. breyeri*. MaxEnt output is the Complementary Log-Log (cloglog) format which is a probability of presence. Mean prediction of 12 model replicates (boostrap) are computed. The mean maximum sensitivity plus specificity (MaxTSS) Cloglog threshold was used to depict in grey values below it (0.61 in the study). Numbers in the map refer to Id number of Table 1.

**S9 Figure.**
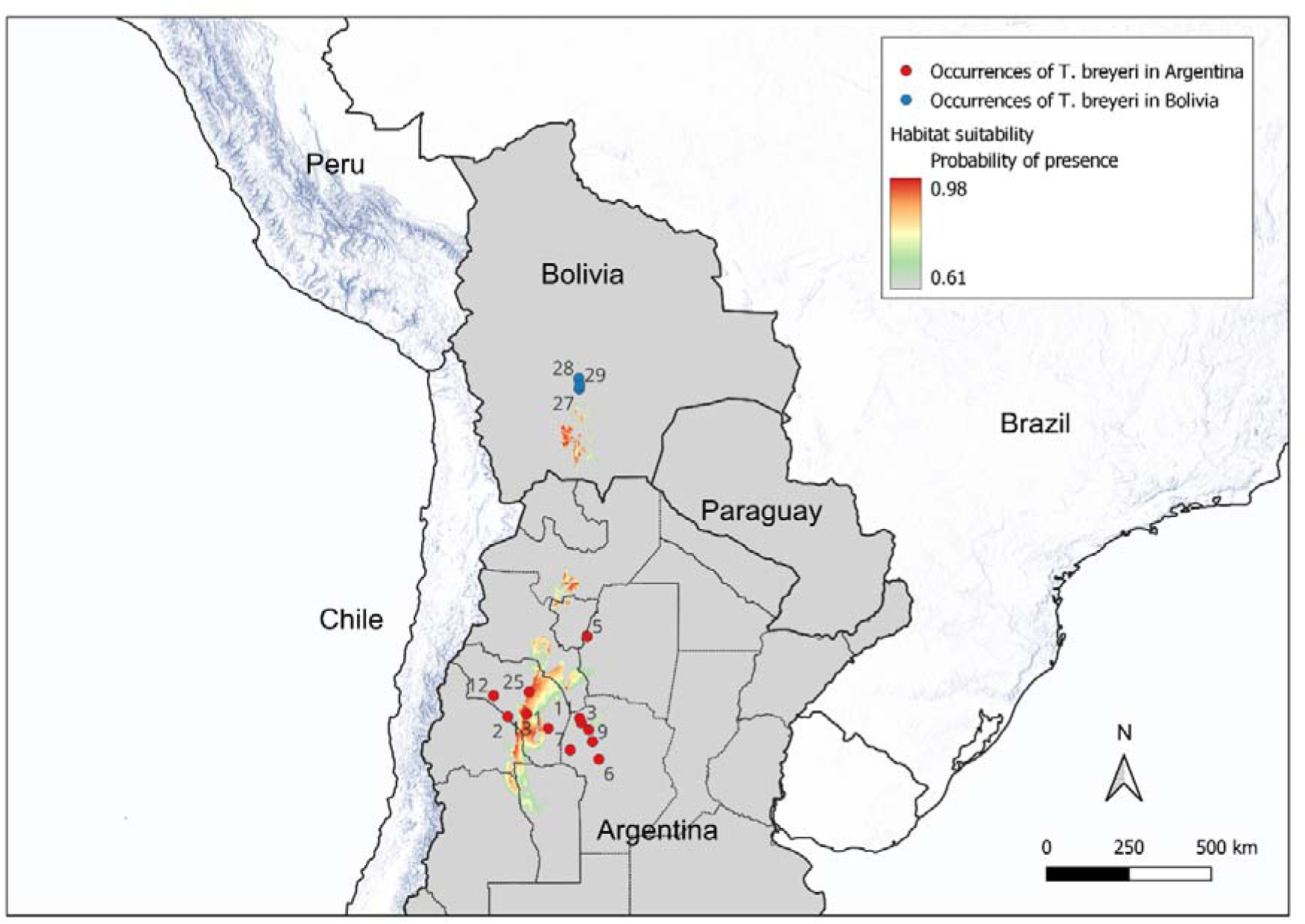
MaxEnt representation of habitat suitability for *T. breyeri*. MaxEnt output is the Complementary Log-Log (cloglog) format which is a probability of presence. Minimum prediction of 12 model replicates (boostrap) are computed. The mean maximum sensitivity plus specificity (MaxTSS) Cloglog threshold was used to depict in grey values below it (0.63 in the study). Numbers in the map refer to Id number of Table 1.

**S10 Figure.**
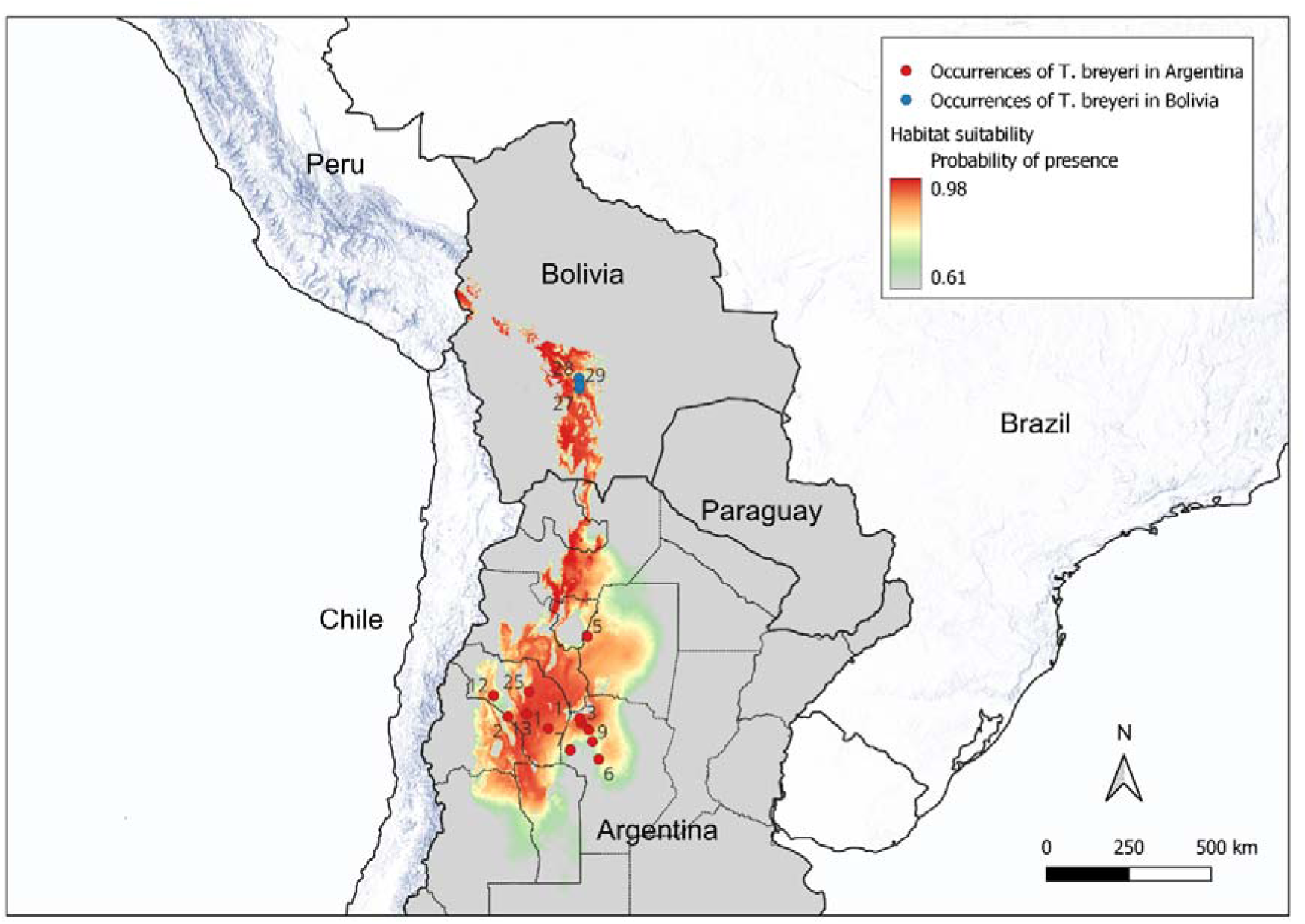
MaxEnt representation of habitat suitability for *T. breyeri*. MaxEnt output is the Complementary Log-Log (cloglog) format which is a probability of presence. Maximum prediction of 12 model replicates (boostrap) are computed. The mean maximum sensitivity plus specificity (MaxTSS) Cloglog threshold was used to depict in grey values below it (0.63 in the study). Numbers in the map refer to Id number of Table 1.

**S1 Table. General outputs of MaxEnt for the selected model**. Each row of the table indicates results for a model from the bootstrap procedure.

**S2 Table.** Bioclimatic variables selected to construct the habitat suitability model with MaxEnt. VIF values are from the step procedure carried out with the *vifstep* function of the R-package *sdm,* with threshold = 5. Correlations are values at the end of the *vifcor* procedure carried out with the *sdm* R-package with threshold = 0.7.

|  |  |  | Correlations |  |  |  |
| --- | --- | --- | --- | --- | --- | --- |
| Variable | Definition | VIF | BIO2 | BIO4 | BIO6 | BIO13 |
| BIO2 | Mean Diurnal Range<br>(Mean of monthly (max temp - min temp)) | 3.16 |  |  |  |  |
| BIO4 | Temperature Seasonality<br>(Standard deviation ×100) | 3.10 | 0.40 |  |  |  |
| BIO6 | Min Temperature of Coldest Month | 3.45 | -0.58 | 0.27 |  |  |
| BIO13 | Precipitation of Wettest Month | 1.91 | -0.60 | -0.35 | 0.45 |  |
| BIO17 | Precipitation of Driest Quarter | 2.06 | -0.56 | -0.54 | 0.27 | 0.53 |

